# Use of convalescent serum reduces severity of COVID-19 in nonhuman primates

**DOI:** 10.1101/2020.10.14.340091

**Authors:** Robert W. Cross, Abhishek N. Prasad, Viktoriya Borisevich, Courtney Woolsey, Krystle N. Agans, Daniel J. Deer, Natalie S. Dobias, Joan B. Geisbert, Karla A. Fenton, Thomas W. Geisbert

## Abstract

Passive transfer of convalescent plasma or serum is a time-honored strategy for treating infectious diseases. Human convalescent plasma containing antibodies against SARS-CoV-2 is currently being used to treat COVID-19 patients. However, most patients have been treated outside of randomized clinical trials making it difficult to determine the efficacy of this approach. Here, we assessed the efficacy of convalescent sera in a newly developed African green monkey model of COVID-19. Groups of SARS-CoV-2-infected animals were treated with pooled convalescent sera containing either high or low to moderate anti-SARS-CoV-2 neutralizing antibody titers. Differences in viral load and disease pathology were minimal between monkeys that received the lower titer convalescent sera and untreated controls. However, and importantly, lower levels of SARS-CoV-2 in respiratory compartments, reduced gross and histopathological lesion severity in the lungs, and reductions in several parameters associated with coagulation and inflammatory processes were observed in monkeys that received convalescent sera versus untreated controls. Our data support human studies suggesting that convalescent plasma therapy is an effective strategy if donors with high level of antibodies against SARS-CoV-2 are employed and if recipients are at an early stage of disease.

---

Severe acute respiratory syndrome coronavirus 2 (SARS-CoV-2) can cause a severe, potentially life threating viral pneumonia named coronavirus disease 2019 (COVID-19) (World.Health.Organisation, 2020). The COVID-19 pandemic originated in Wuhan, China and spread across the globe at an explosive rate leading to over 21 million cases and hundreds of thousands of deaths in just over 6 months’ time. While no currently approved vaccines or therapeutics exist for COVID-19, a battery of medical countermeasures are being developed and assessed in human clinical trials at record speed (Beigel et al., 2020; Jackson et al., 2020), (Zhu et al., 2020). Nonetheless, safety and efficacy trials can take considerable time to produce confidence prior to obtaining regulatory approvals, which creates an immediate need for therapeutic options that may be more accessible; ideally one with track records of success against related viruses. Transfusion of convalescent blood products has been used in clinical settings for >100 years (Casadevall et al., 2004) including for the treatment of emerging viruses such as Ebola, influenza, and other viruses (Mair-Jenkins et al., 2015).

Immunity to the closely related severe acute respiratory syndrome coronavirus (SARS-CoV) and Middle East respiratory syndrome coronavirus (MERS-CoV) is understood to be owed, in part, to the development of potent neutralizing antibody responses (Sariol and Perlman, 2020). Indeed, some of the first approaches to treat humans for acute cases that were otherwise unresponsive to standard respiratory virus treatment protocols was the administration of convalescent plasma (CP) from SARS-CoV (Yeh et al., 2005) or MERS-CoV (Arabi et al., 2015), respectively. Recently, the World Health Organization approved a standardized protocol for the use of CP for the treatment of MERS-CoV (Arabi et al., 2015), yet there is still debate on the feasibility of CP use in the treatment of MERS as donor antibody titers tend to be too low to produce therapeutic effect (Corti et al., 2016). Nonetheless, with ever increasing caseloads and no other immediately available options, reports of the use of COVID-19 convalescent plasma (CCP) for the treatment of severe COVID-19 patients surfaced which suggest therapeutic benefit even if given in some cases of severe disease (Joyner et al., 2020a; Shen et al., 2020). Building on these successes, large scale clinical trials have been initiated in association with nationwide donation programs (CCPP19, 2020; Malani et al., 2020). In response to the worsening public health crisis, the United States Food and Drug Administration issued a federal emergency use authorization (EUA) for the use of CCP on 23 August, 2020 to facilitate access to the treatment approach, despite ongoing clinical trials to fully evaluate safety and efficacy (US-FDA, 2020).

In parallel with reports of success in humans, hamsters were recently used to experimentally demonstrate the potential of CCP to treat SARS-CoV-2 infection. While informative to demonstrate proof-of-concept, limited reagents are available to succinctly describe the host responses to infection and treatment in hamsters (Chan et al., 2020; Imai et al., 2020). Non-human primates (NHP) have long been used to model pathological responses to infection due to their physiological similarity to humans and abundance of cross-reactive reagents, which allow for more detailed analysis than possible in lower vertebrates. A natural extension to this utility is their value in determining predictive efficacy of medical countermeasures such as vaccines or therapies in humans. Several groups have recently described NHP infection in a number of species including rhesus macaques, cynomolgus macaques, baboons, and marmosets (Lu et al., 2020; Rockx et al., 2020; Singh et al., 2020), none of which elicit overt clinical signs of disease reflecting the human condition, making evaluation of therapeutic approaches in these species challenging. Recently we described a novel African green monkey (AGM) model which recapitulates many of the most salient features of human disease including severe viral pneumonia, transient coagulopathy, and a prolonged state of recovery (Cross et al., 2020; Woolsey et al., 2020). In this study, we challenged AGMs with SARS-CoV-2 and subsequently treated the animals with different pools of sera derived from AGM previously infected with the virus. We provide direct evidence for the importance of neutralization potency on the efficacy of convalescent sera to reduce viral burden in the respiratory tract and to reduce systemic and localized evidence of COVID-19 in AGMs.

## Results

### SARS-CoV-2 infection of African green monkeys and passive treatment with convalescent serum

Ten healthy, SARS-CoV-2 naïve AGMs were randomized into two treatment cohorts (n=4 each) and an untreated control cohort (n=2). All animals were challenged with a target dose of 5.0 x 10^5^ PFU of SARS-CoV-2 (SARS-CoV-2/INMI1-Isolate/2020/Italy) via combined intranasal (i.n.) and intratracheal (i.t.) inoculation. Ten hours post-challenge, the experimental cohorts were treated intravenously (i.v.) with pooled convalescent sera (6.1 ml/kg) obtained from animals infected with the homologous isolate of SARS-CoV-2 in previous studies (Cross et al., 2020; Woolsey et al., 2020). Animals in one experimental cohort received pooled sera from a group of three AGMs back-challenged 35 days after primary challenge, and then euthanized at the scheduled study endpoint 22 days after the back-challenge. To determine the SARS-CoV-2 binding and neutralizing potential of the treatment *in vivo*, sera from the convalescent sera-treated AGMs was fractionated from whole blood collected 2 and 5 dpi and assessed for binding by ELISA and neutralizing activity by PRNT_50_ (**Figure 1a,b**). The binding titer was 1:51,200 (total virus), 1:25,600 (anti-Nucleoprotein), and 1:12,800 (anti-spike RBD); which corresponded to a PRNT_50_ in pooled sera of ~ 1:2048 for this cohort (designated “high dose”; “HD”). The second experimental cohort received pooled sera from a group of three AGMs euthanized at the scheduled study endpoint 34 days after challenge with SARS-CoV-2 (Cross et al., 2020); the binding titer was 1:3,200 (total virus, anti-Nucleoprotein, and anti-spike RBD); which corresponded to a PRNT_50_ for pooled sera of ~ 1:128 (designated as “low dose”; “LD”). Animals were longitudinally monitored for clinical signs of illness and euthanized 5 dpi.

**Figure 1:**
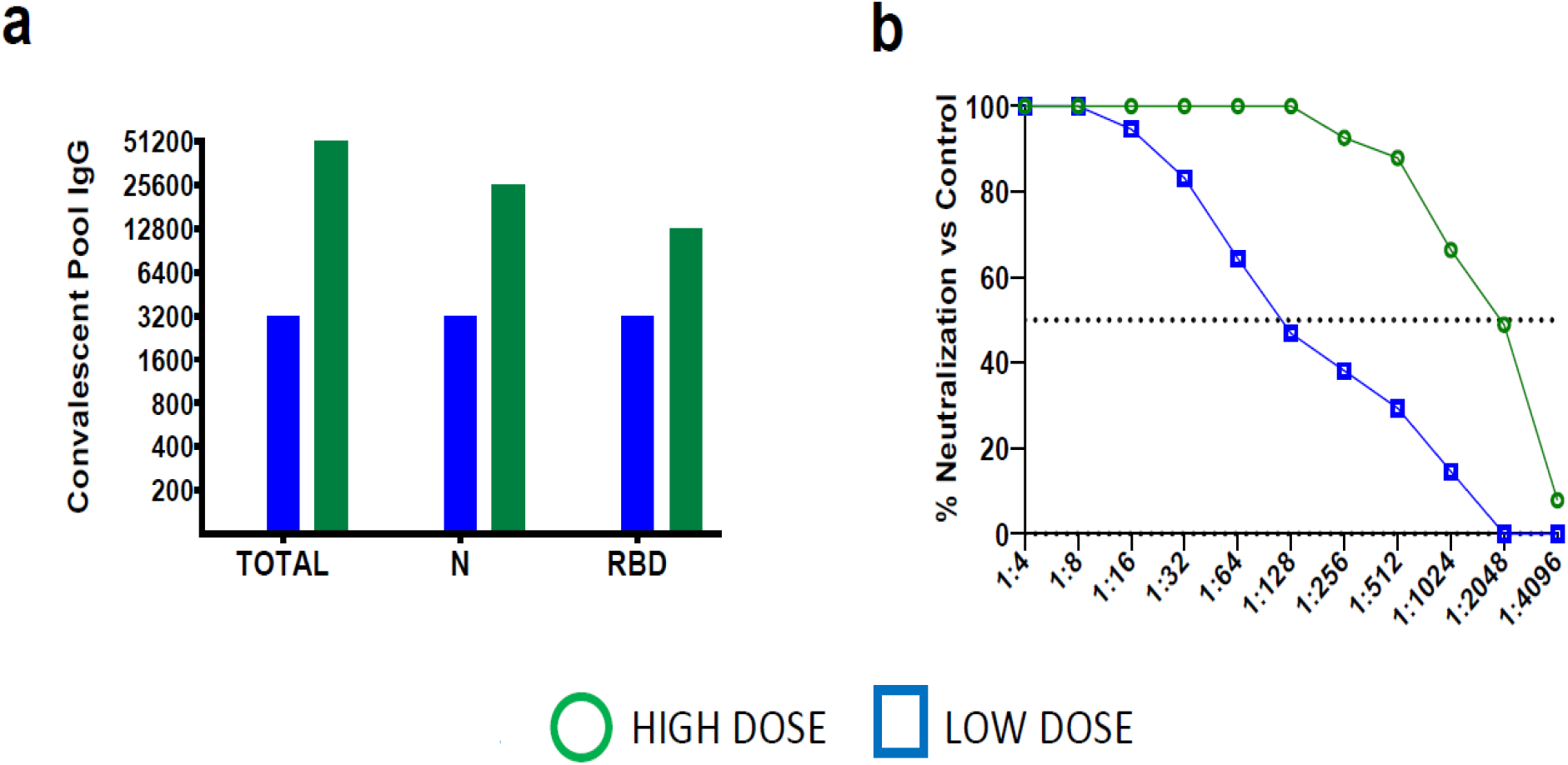
Characterization of convalescent COVID19 plasma pools from SARS-CoV2 infected African Green Monkeys: (a.) Pooled convalescent plasma was assayed by ELISA for antibodies reactive to SARS-CoV2 virus cellular lysates (Total), nucleoprotein (N), or spike protein receptor binding domain (RBD). (b.) PRNT_50_ assays were performed on pooled convalescent sera from AGMs challenged with the homologous isolate of SARS-CoV-2 in previous studies (Cross et al., 2020; Woolsey et al., 2020).

### Clinical disease

In agreement with previous reports (Cross et al., 2020; Hartman et al., 2020; Johnston et al., 2020; Speranza et al., 2020; Woolsey et al., 2020), SARS-CoV-2-infected AGMs in this study showed mild to moderate clinical illness (**Table 1**). Shifts in leukocyte populations as compared to pre-challenge baseline counts; specifically, lymphocytopenia, generalized granulocytopenia (neutropenia, eosinopenia, and/or basopenia), and mild to moderate thrombocytopenia were common to most animals regardless of cohort starting approximately 2 dpi (**Table 1**). Four animals (HD-AGM-3, LD-AGM-3, LD-AGM-4, and C-AGM-2) experienced monocytosis beginning 2-4 dpi; in two of these animals (HD-AGM-3 and LD-AGM-4) this coincided with generalized granulocytosis (neutrophilia, eosinophilia, and/or basophilia). Prothrombin time (PT) was largely unaffected, yet a significantly prolonged coagulation time was noted at 4 dpi for activated partial thromboplastin time (aPTT) in HD versus control group (p=0.0459; two-way ANOVA with Tukey’s multiple comparison); decreases in levels of thrombocytes were more notable in control animals; and increases in circulating fibrinogen were generally more pronounced in the control group compared to either of the experimental groups (**Figure S1**). These results suggest that while all animals appear to have experienced varying degrees of coagulopathy, treatment with sera with a higher SARS-CoV-2 neutralizing capacity may have partially ameliorated disease in the HD group. However, differences in these parameters were not statistically significant for most time points due to the small cohort sizes and individual animal variability. Serum markers of renal and/or hepatic function (CRE, ALT, AST) were mildly elevated in most animals from treatment groups as well as the control group, while CRP, a marker of acute systemic inflammation, was mild to moderately (1-10 fold over baseline) elevated 4-5 dpi in all but a single animal (LD-AGM-2).

**Table 1.**
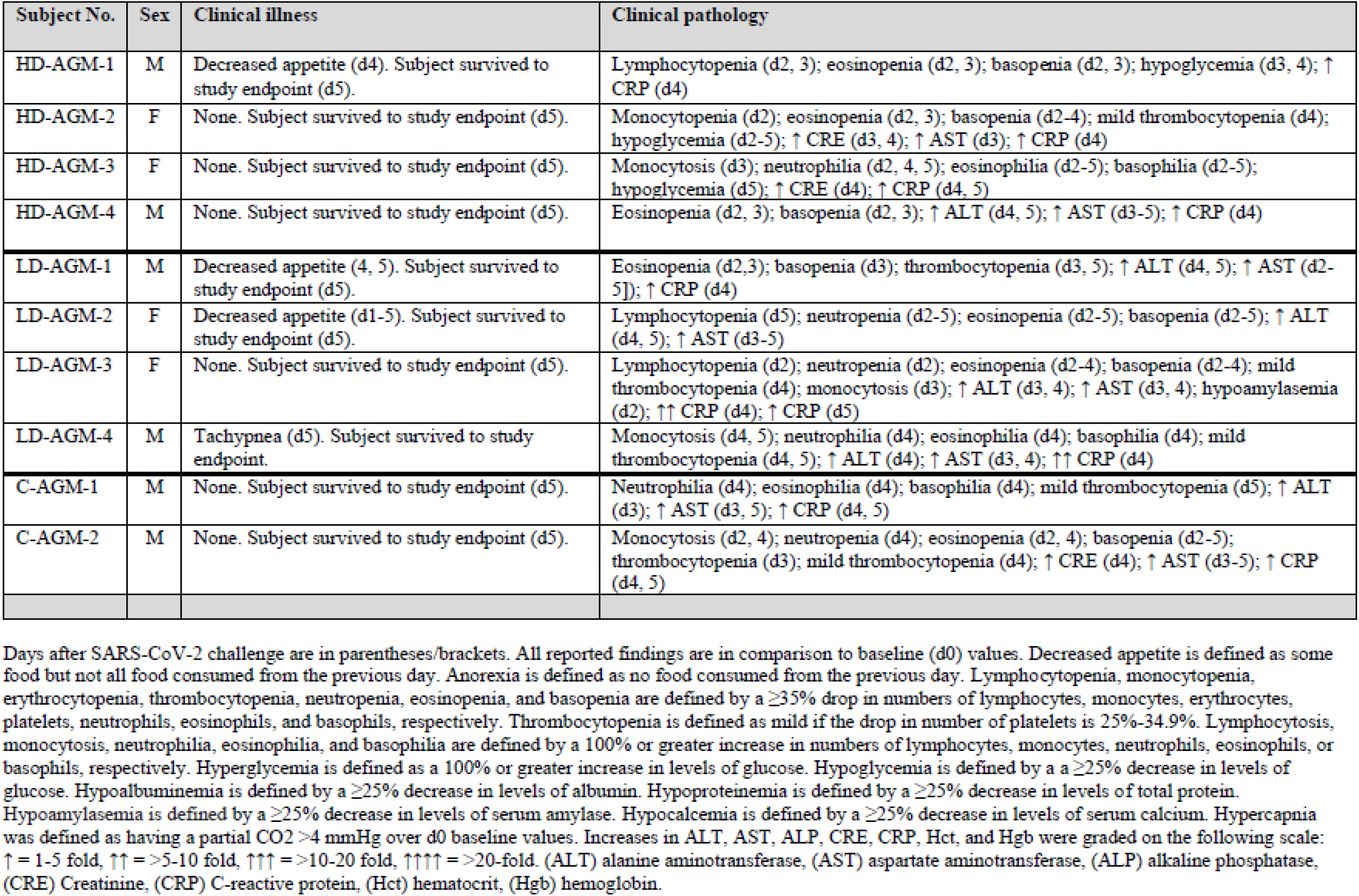
Clinical description and outcome of African green monkeys following SARS-CoV-2 challenge.

### Assessment of antibody levels

Circulating SARS-CoV-2 specific antibodies to total virus, nucleoprotein, and the receptor binding domain (RBD) of the spike protein were higher in all animals on 2 dpi of the HD group compared to the LD group and untreated subjects, but by day 5, spike RBD titers in 2/4 HD animals began to decline slightly (**Figure 2a,b,c**). Three of four animals in the HD-treated group had mean PRNT50 titers of ~ 1:128 at 2 dpi, while the fourth (HD-AGM-1) had a neutralizing titer between 1:64 and 1:128 (**Figure 2d**). Neutralizing titers in AGMs that have survived experimental SARS-CoV-2 challenge have previously been demonstrated to be as low as 1:16, suggesting low neutralizing titers may be negligible when considered as a part of the entire immune response (Cross et al., 2020; Woolsey et al., 2020). Neutralizing antibody titers were markedly lower in LD-treated animals and a nearly complete lack of neutralizing activity was observed in the untreated control animals at 2 dpi. At 5 dpi, neutralizing antibody titers waned to ~ 1:64 in HD-treated animals and between 1:4 and 1:8 in LD-treated animals. A single control animal (C-AGM-1) had similar neutralizing activity at this time point (**Figure 2e**). The rapid decrease in circulating neutralizing antibody titers by 5 dpi and the lack of robust neutralization in the control and LD-treated animals indicated that much of the neutralizing activity in the HD-treated group was acquired through treatment.

**Figure 2:**
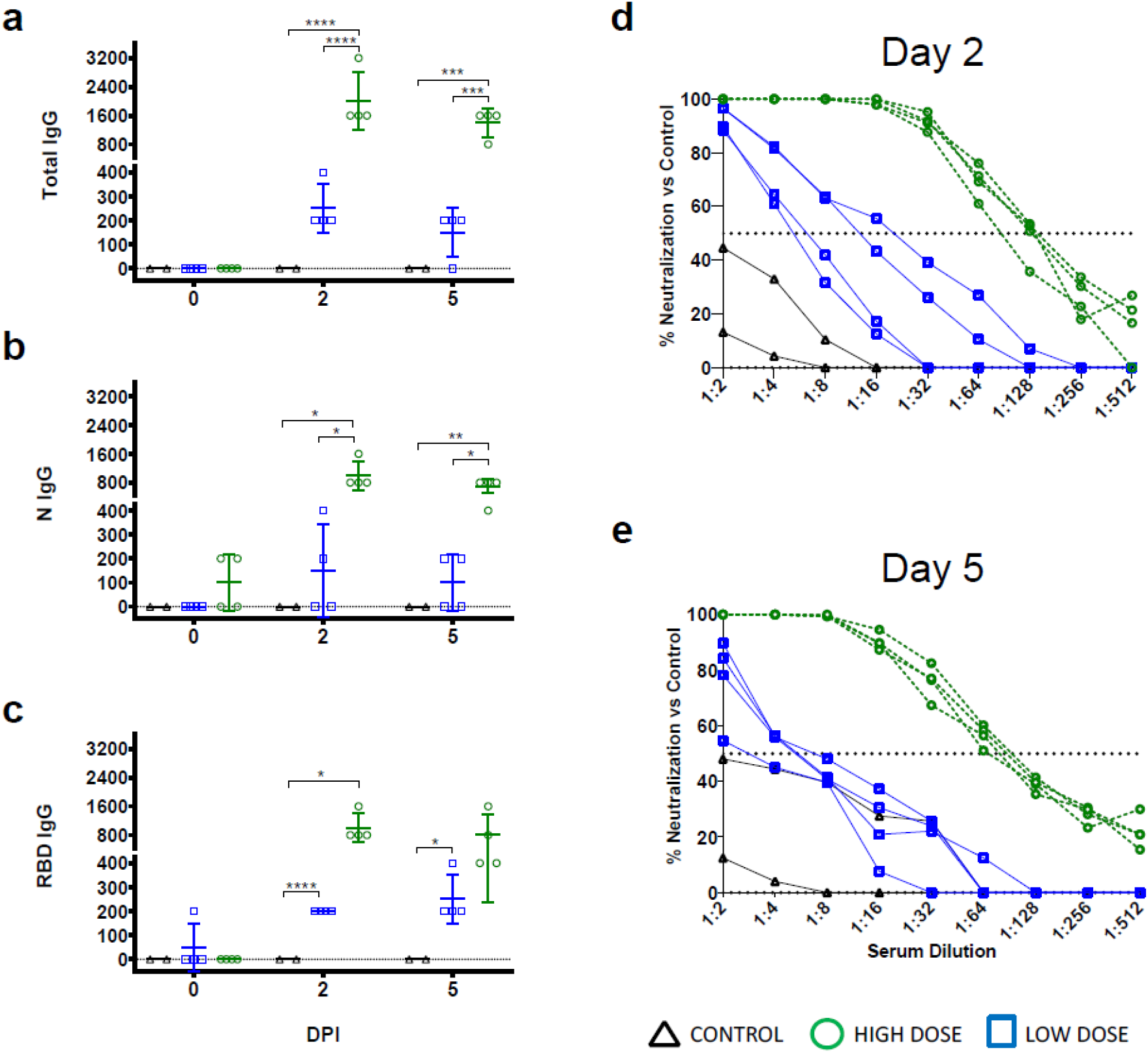
Serology of SARS-CoV-2-challenged AGMs treated with convalescent sera. ELISA binding titers of sera collected on the indicated time points from AGMs enrolled in the current study following challenge with SARS-CoV-2 and passive transfer of convalescent sera for IgG against SARS-CoV2 virus cellular lysates (Total, (a)), nucleoprotein (N, (b)), or spike protein receptor binding domain (RBD, (c)). PRNT50 assays were performed on pooled convalescent sera from AGMs challenged with the homologous isolate of SARS-CoV-2 in previous studies (Cross et al., 2020; Woolsey et al., 2020) compared with control animals on day 2 post infection (d) and day 5 post infection (e). Data shown is the percent reduction in SARS-CoV-2 plaque counts (mean of duplicate wells) compared to a control plate (no sera).

### Assessment of viral load

We next assessed viral load in whole blood and mucosal swabs on 0, 2, 3, 4, and 5 dpi, bronchoalveolar lavage (BAL) fluid on −7, 3, and 5 dpi, and lung homogenates at the study endpoint (5 dpi) by both RT-qPCR detection of viral RNA (vRNA) and plaque titration of infectious virus. As previously reported (Cross et al., 2020; Woolsey et al., 2020), there was no circulating SARS-CoV-2 detected in the peripheral blood, as assessed by RT-qPCR of whole blood and plaque titration of the plasma fraction, respectively (data not shown). vRNA was detected in nasal swabs from three animals from the HD-treated group, four animals from the LD-treated group, and all animals from the control group, including 6/6 historical controls (**Figure 3a**). Notably, in two animals from the HD-treated group, detection of vRNA was limited to a single day (2 dpi in HD-AGM-2 and 5 dpi in HD-AGM-3). A low amount (~ 2.4 log10 PFU/mL) of infectious SARS-CoV-2 was recovered from nasal mucosa in a single animal from the HD-treated group 2 dpi, while similar amounts of virus were detected on multiple time points for most animals from both the LD-treated and control groups (**Figure 3e**). Interestingly, while levels of vRNA and infectious virus from the oral mucosa were generally lower and less frequently detected in HD-treated animals compared to LD-treated animals and the two control animals from this study, both vRNA and infectious virus were only detected from 3/6 animals from historical controls, despite animals from both studies being challenged with identical virus stock via identical challenge route (**Figure 3b,f**). Detection of vRNA from the rectal mucosa was similarly variable, present in samples from only 2/4 animals from the HD-treated group, 3/4 from the LD-treated group, and only two control animals (C-AGM-2 and a single HC-AGM), while infectious virus was not recovered from the rectal swabs of any animal (**Figure 3d, and data not shown**). Strikingly, while SARS-CoV-2 vRNA was detected at similar levels in the BAL fluid from all animals at 3 dpi, only 2/4 animals in the HD-treated group had detectable vRNA at 5 dpi, compared to all of the animals in both the LD-treated and control groups (**Figure 2c**), while infectious virus was not recoverable at all from the BAL fluid of both treated groups on day 5. Conversely, infectious virus was detected from both controls from this study and all historical controls (**Figure 3g**). Genomic vRNA was detected in similar quantities from the lower, middle, and upper sections of the lungs from all animals in all groups (**Figure 3h**). While comparisons of viral load in mucosal compartments failed to reach statistical significance, taken together, these data suggest that treatment with convalescent sera may decrease viral replication and shedding and thus mitigate disease and transmission.

**Figure 3:**
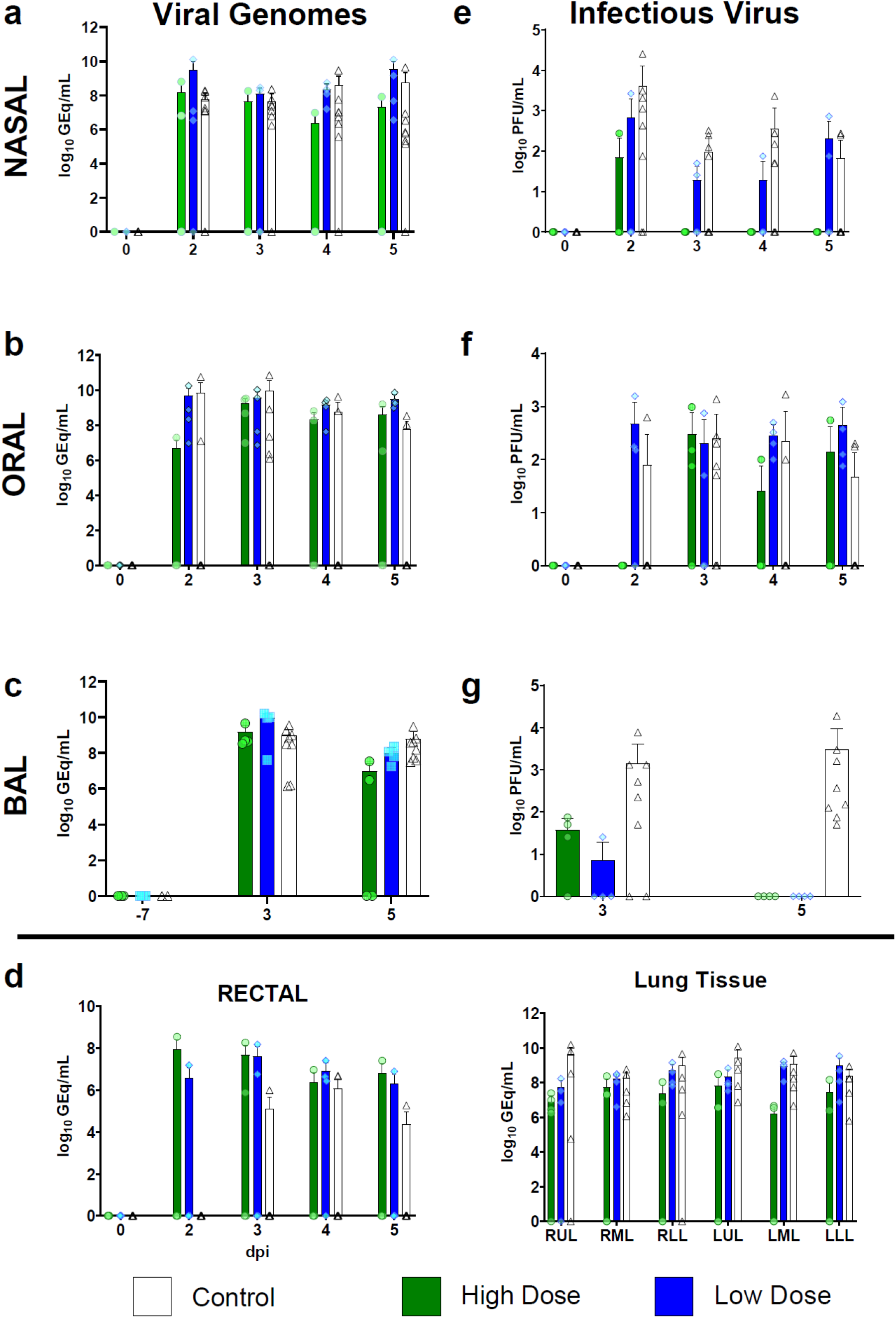
Viral load in mucosal swabs, BAL fluid, and lung tissue from SARS-CoV-2-infected AGMs. SARS-CoV-2 viral load was assessed by plaque titration (**B, D, F**) and/or RT-qPCR (**A, C, E, G, H**) from mucosal swabs and BAL fluid collected at the indicated timepoints, and lung tissue harvested at necropsy (5 dpi). For all panels, individual data points represent the mean of two technical replicates from a single assay. Dashed horizontal lines indicate the limit of detection (LOD) for the assay (1000 GEq/mL for RT-qPCR; 25 PFU/mL for plaque titration). Bars indicate the mean for each cohort at the indicated time point, values for individual animals within each cohort are shown as color-coded symbols. Historical control animals from a previous study (Woolsey et al., 2020) utilizing the homologous virus are included for statistical purposes. Error bars indicate the upper bound SD. To fit on a log scale axis, zero values (below LOD) are plotted as “1” (10^0^); however, statistical comparisons were performed using the original zero values. Statistical significance was assessed by two-way ANOVA with the Geisser-Greenhouse correction without the assumption of sphericity, except for **(E)** where mixed-effects modeling was used to account for missing historical control values at −7 dpi, and **(F)** where no correction for sphericity was necessary as only two time points were being compared. Tukey’s post-hoc test for multiple comparisons was used to assess differences between cohorts at matched time points. Statistically significant comparisons are indicated by asterisk(s) (*= 0.05 ≤ p ≤ 0.01, **= 0.01 ≤ p ≤ 0.001).

### Comparative Pathology

On 5 dpi, all animals were euthanized to gauge viral burden and determine pathological changes in the lungs associated with the treated infection. Multifocal pulmonary consolidation with hyperemia and hemorrhage were noted in all AGMs at 5 dpi. In all AGMs, the most severe lesions were located in the dorsal aspects of the lower lung lobes (**Figure 4**). A board-certified veterinary pathologist approximated gross lesion severity for each lung lobe. Gross lung scores were most severe in untreated control AGMs followed by the LD-treated animals and least severe among the HD-treated AGMs (**Figure 4, S2**). Mild lymphoid enlargement was noted in one untreated control AGM and two LD-treated AGMs.

**Figure 4:**
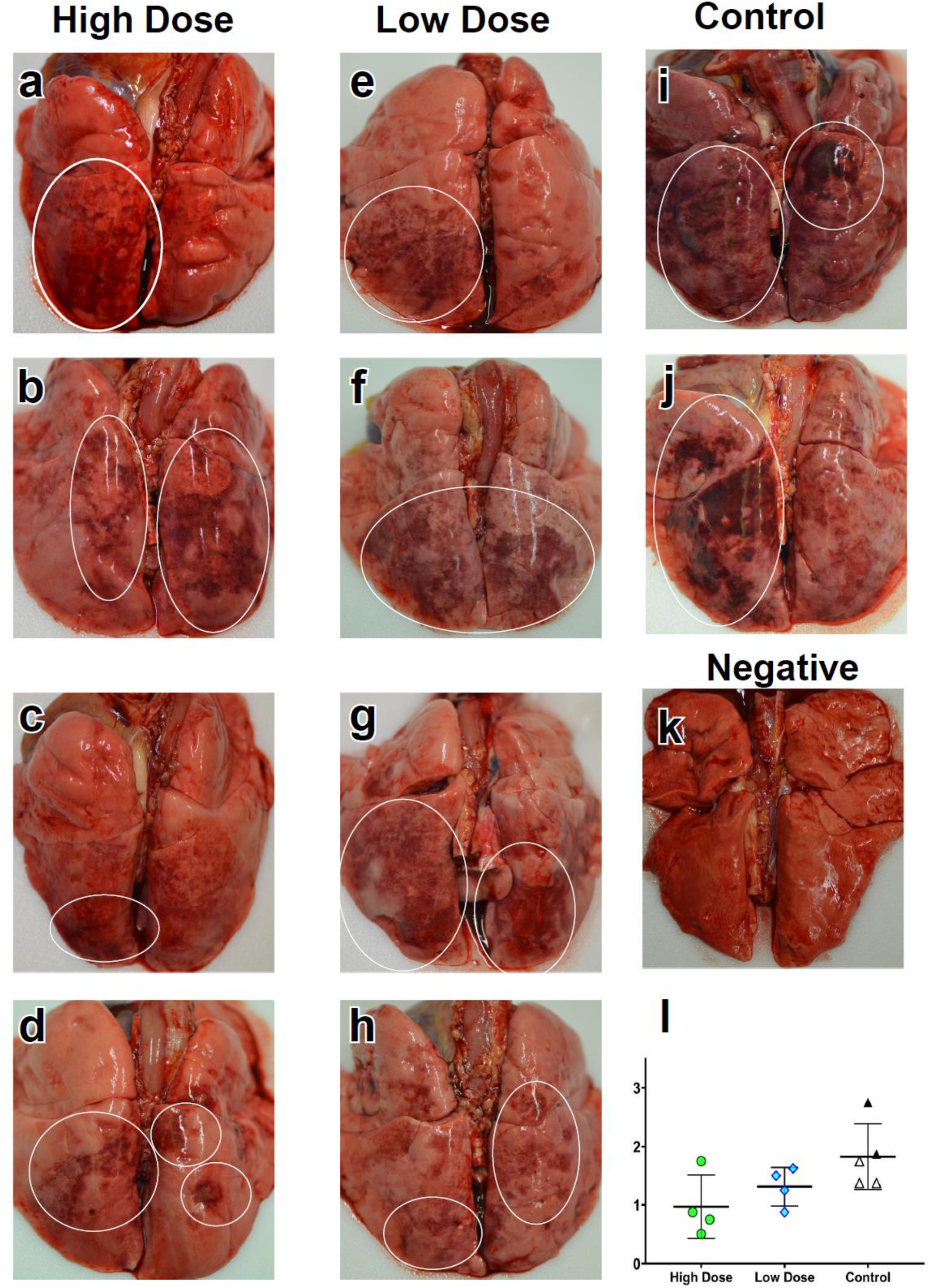
Dorsal view of gross pulmonary lesions in AGMs infected with SARS-CoV-2 with and without treatment. Lungs from AGMs treated with high neutralizing antibody titers (HD-AGM-1 (a), HD-AGM-2(b), HD-AGM-3(c), HD-AGM-4(d)) fail to collapse and present with mild locally extensive pulmonary consolidation with hyperemia and hemorrhage (circled regions). Lungs from AGMs treated with low neutralizing antibody titers (LD-AGM-1(e), LD-AGM-2(f), LD-AGM-3(g), & LD-AGM-4 (h)) fail to collapse and present with moderate locally extensive pulmonary consolidation with hyperemia and hemorrhage (circled regions). Lungs from positive control AGMs (C-AGM-1 (i) & C-AGM-2(j)) euthanized 5 dpi after infection with SARS-CoV-2 fail to collapse and present with marked locally extensive pulmonary consolidation with hyperemia and hemorrhage (circled regions). Dorsal view of control lungs with no significant lesions from SARS-CoV-2 negative AGMs (l) Gross examination severity scores by group (m).

Histologically, the untreated control AGMs euthanized at 5 dpi developed interstitial pneumonia and multifocal alveolar flooding with edema, fibrin, red blood cells and mixed inflammatory cells, as previously observed (Cross et al., 2020; Woolsey et al., 2020). The interstitial pneumonia was characterized by multifocal moderate expansion of alveolar septae with macrophages, lymphocytes, and fewer numbers of neutrophils (**Figure 5a**). Bronchial respiratory epithelium was multifocally ulcerated with associated underlying acute inflammation. Modest expansion of alveolar septae with collagen was noted multifocally (**Figure 5d**). Colocalization of SARS-CoV-2 antigen with pulmonary lesions were demonstrated with antibodies against SARS-CoV-2 N protein. Positive immunohistochemical labeling was noted diffusely within the cytoplasm of respiratory epithelium of the bronchus, type I pneumocytes, type II pneumocytes and rarely alveolar macrophages (**Figure 5b**).

**Figure 5:**
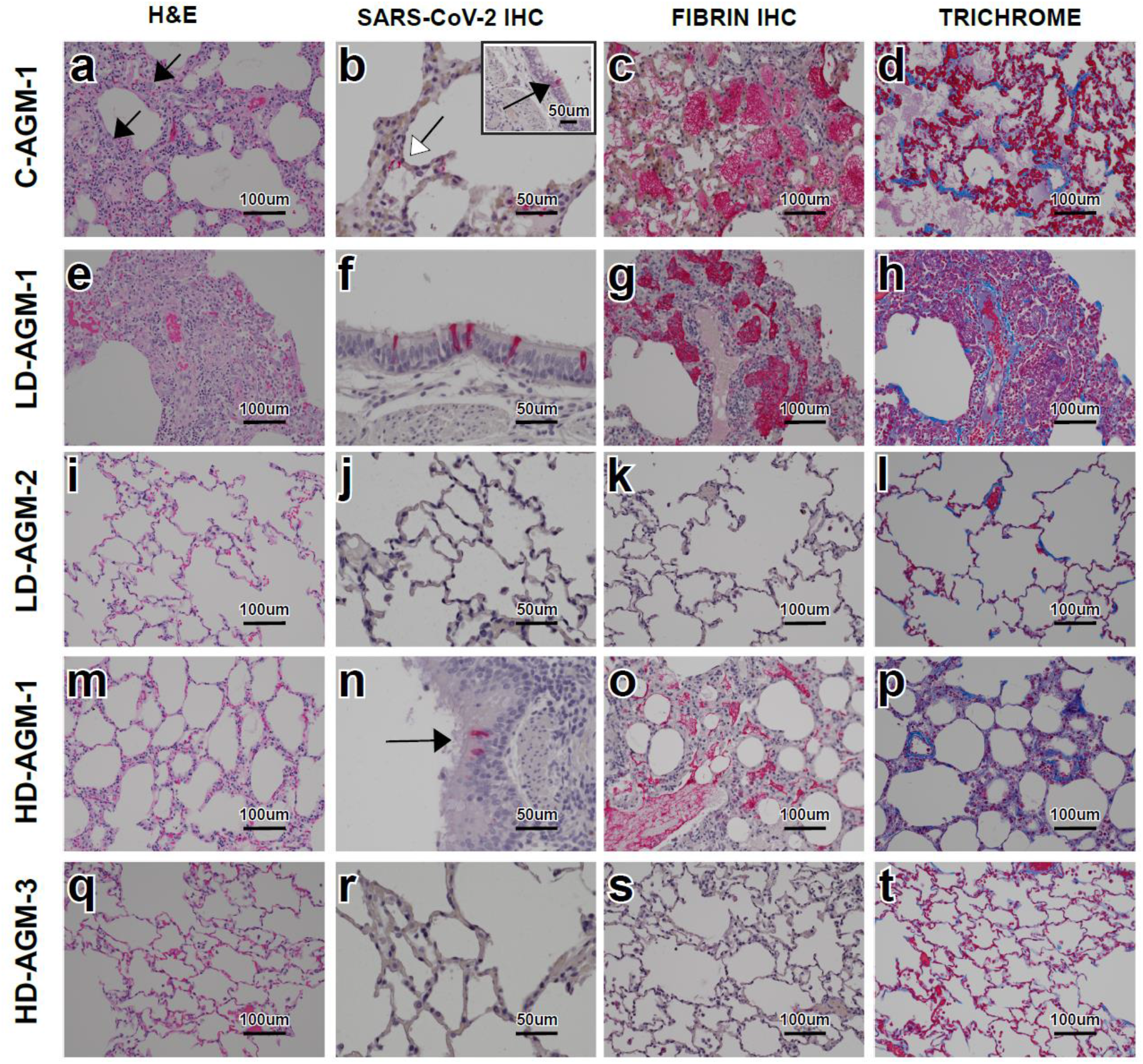
Pulmonary histologic changes in AGMs infected with SARS-CoV-2 with and without treatment. (A-D) Lung lobe of positive control C-AGM-2(A) Multifocal interstitial pneumonia with multinucleated cells (black arrow), (B) SARS-CoV-2 IHC positive (red) pneumoncytes (white arrows) and bronchial respiratory epithelium (inset, black arrow), (C) Flooded alveolar spaces with fibrin (red), (D) Modest collagen deposition (blue) in alveolar septum. (E-H) Lung lobe of low neutralizing antibody titer treatment LD-AGM-1 (E) Multifocal interstitial pneumonia, (F) SARS-CoV-2 IHC positive (red) bronchial respiratory epithelium, (G) Flooded alveolar spaces with fibrin (red), (H) Modest collagen deposition (blue) in alveolar septum. (I-L) Lung lobe of low neutralizing antibody titer treatment LD-AGM-2 (I) No significant lesions, (J) No immunolabeling for anti-SARS-CoV-2 antigen (K) No immunolabeling for anti-fibrin antigen (L) No significant findings. (M-P) Lung lobe of high neutralizing antibody titer treatment HD-AGM-1 (M) Mild multifocal interstitial pneumonia, (N) SARS-CoV-2 IHC positive (red) bronchial respiratory epithelium, (O) partially flooded alveolar spaces with fibrin (red), (P) Modest collagen deposition (blue) in alveolar septum. (Q-T) Lung lobe of high neutralizing antibody titer treatment HD-AGM-3 (Q) No significant lesions, (R) No immunolabeling for anti-SARS-CoV-2 antigen (S) No immunolabeling for anti-fibrin antigen (T) No significant findings. H&E staining 20x (A, E, I, M, & Q), IHC labeling for anti-SARS-CoV2 antigen 40x (red) (B, B inset, F, J, N & R), IHC labeling for anti-fibrin antigen 20x (red) (C, G, K, O & S), Trichrome staining 20x (blue) (D, H, L, P & T).

The eight AGMs treated with convalescent sera developed varying degrees of multifocal pulmonary lesions. Five treated AGMs, 3/4 from the LD cohort and 2/4 from the HD cohort developed moderate pulmonary lesions, similar to the untreated control AGMs (**Figure 5e, g, h, m, o, p**). SARS-CoV-2 antigen detection in these AGMs was noted diffusely within the cytoplasm of individual to small clusters of respiratory epithelium of the bronchus (**Figure 5f & n**). The remaining three treated AGMs, one from the LD cohort (LD-AGM-2) and two from the HD cohort (HD-AGM-3 & HD-AGM-2), had minimal multifocal interstitial pneumonia, characterized by minimal expansion of alveolar septae with macrophages, lymphocytes and rarely neutrophils (**Figure 5 i & q**). There was no positive immunolabeling for SARS-CoV-2 in these animals, minimal to no fibrin flooding of alveolar spaces and no overt expansion of alveolar septae with collagen (**Figure 5 j-l, r-t**).

### Assessment of soluble inflammation markers

We measured plasma concentrations of a panel of cytokines/chemokines known to be altered in human COVID-19 (Laing et al., 2020). On 2 dpi, Interleukin-10 (IL-10) and IL-12 were significantly elevated in control animals compared to both LD and HD groups [IL-10: p=0.0047 (HD) and 0.0210 (LD) and IL-12p40: p=<0.0001(HD) and 0.0015 (LD) two way ANOVA supported by Tukey’s multiple comparison test] (**Figure 6b, e**). Interferon (IFN) gamma-induced protein 10 (IP-10) was significantly elevated only in the LD cohort [p=<0.0001 (compared to control) and <0.0001 (compared to HD) two way ANOVA supported by Tukey’s multiple comparison test] (**Figure 6c**). Significant elevations in macrophage chemotactic protein 1 (MCP-1) were observed in the all study and historical control animals [p=0.0109 (compared to HD) two way ANOVA supported by Tukey’s multiple comparison test]as well as the LD [p=0.0006 (compared to LD) two way ANOVA supported by Tukey’s multiple comparison test] group on day 2 (**Figure 6k**). On day 3, IL-6 was elevated only in two control animals. (**Figure 6a**). Reduced levels of IFN beta, tumor necrosis factor (TNF) alpha, IFN gamma, IL-8, and MCP-1 were observed in all of the HD-treated animals as compared to control groups beginning 2 dpi and trending through the remainder of the study (**Figure 6g, h, i, j, and k**).

**Figure 6:**
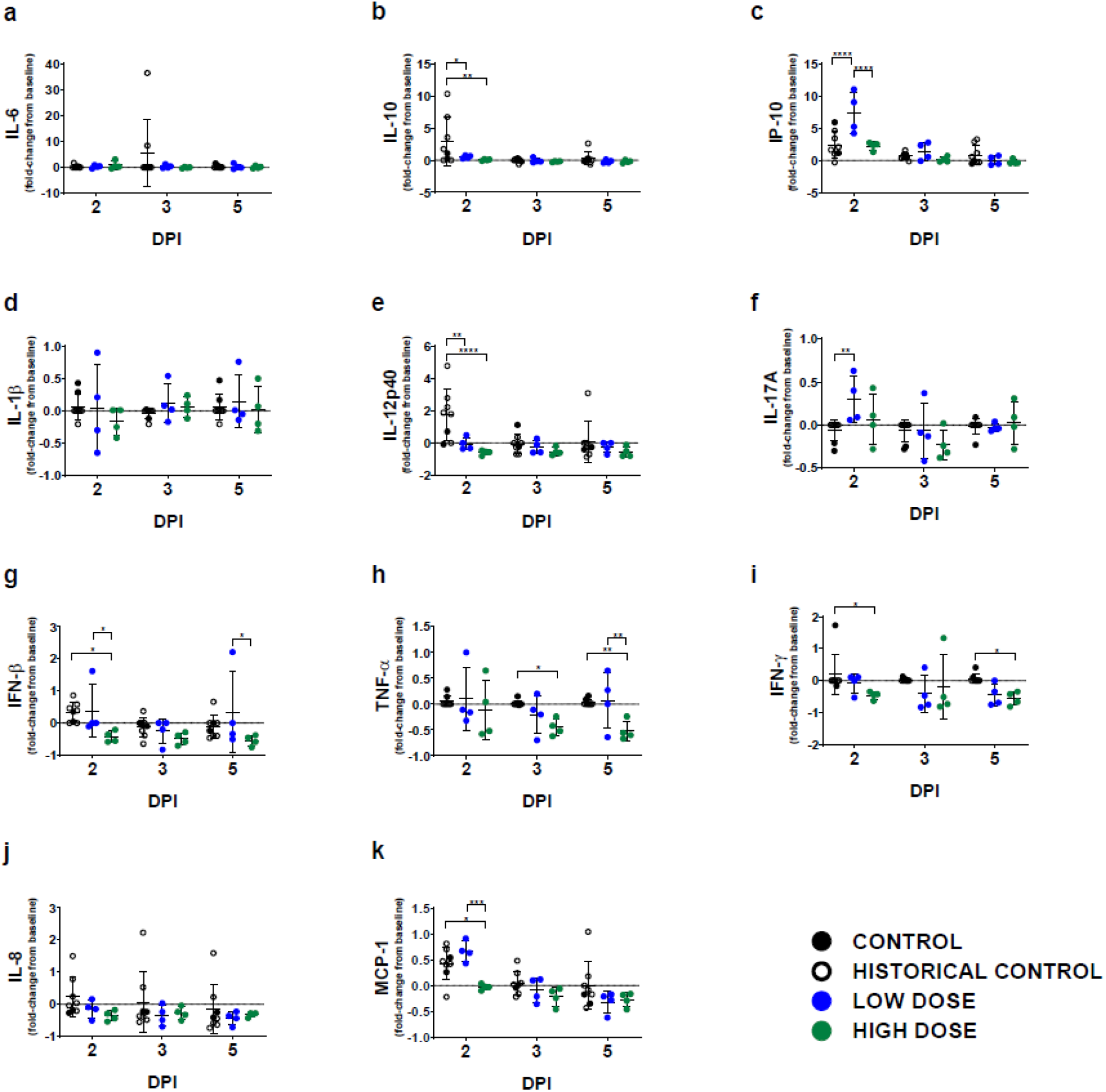
Circulating levels of inflammatory markers. Fold change from individual subject baselines (0 dpi) were calculated for each subject at the indicated timepoints. A two-way ANOVA and Tukey’s multiple comparisons test was used to compare each treatment group at the respective time point (*, p < 0.0332; **, p < 0.0021; ***, p < 0.0002; ****, p < 0.0001). Shown is the average value of duplicate samples for each subject and analyte. Historical control animals from a previous study (Woolsey et al., 2020) utilizing the homologous virus were included for statistical purposes.

## Discussion

In response to the dire need for therapeutic options, several human clinical trials have been initiated to evaluate both the safety and the efficacy of passive transfer of CCP to acutely ill COVID-19 patients (Joyner et al., 2020a). Overall, the consensus of these studies is that therapeutic benefit is possible; however, a number of caveats associated with doing human studies including unknown incubation times, effective dose, and/or route(s) of exposure. These shortcomings highlight the need for experimentally-controlled studies to help understand potential risks of this approach.

Others have demonstrated therapeutic benefit in rodent models of COVID-19 but limited information was gleaned in terms of the pathophysiological parameters impacted by treatment with CCP (Chan et al., 2020; Imai et al., 2020). While hyper-immune plasma has been shown to provide therapeutic benefit in related MERS-CoV-infected primates (van Doremalen et al., 2017), to date, there are no reports demonstrating convalescent sera or plasma treatment benefit in NHP models of COVID-19. Here, we provide evidence that convalescent sera with high neutralizing antibody titers can be given as a postexposure treatment using a NHP model which faithfully recapitulates hallmark features of human COVID-19. Compared to LD-treated or untreated control animals, HD-treated AGMs had lower viral burdens in respiratory compartments, reduced gross and histopathological lesion severity in the lungs, and reductions in several clinical parameters previously shown to be important in human disease including: pro-longed coagulation times, elevated fibrinogen, thrombocytopenia, and hypercytokinemia. Differences in clinical parameters of the LD-treated group with untreated control animals from this study or historical control animals were minimal; however, the lack of infectious SARS-CoV-2 in the BAL samples from all of the LD-treated animals and reduced lung pathology suggest that an antiviral effect was present despite the lower concentration of neutralizing antibodies in the dose of convalescent sera administered. Nonetheless, these data support the need for high potency neutralizing antibody content in convalescent blood product preparations to achieve maximal therapeutic benefit.

Previous work utilizing passive transfer of potent neutralizing antibody containing plasma derived from a cohort of rhesus macaques vaccinated with a SARS-CoV spike expressing ADC-MVA vaccine resulted in exacerbation of pulmonary disease suggestive of antibody dependent enhancement (ADE) (Liu et al., 2019), which served to caution the design of therapeutics and vaccines of potential dangers of targeting this antigen for SARS-CoVs. While we characterized the presence of antibodies capable of binding all SARS-CoV-2 proteins, and more specifically nucleoprotein and the RBD of spike, we did not assess the differential binding potential of all of these targets in the treatments provided. Therefore, we cannot rule out other antiviral mechanisms such as antibody-binding capacity or fragment crystallizable (FC) region-directed functions associated with antibodies present in these treatments. Nor can we account for other soluble factors that may have been present in sera derived from survival of natural infection, which may explain the differences observed in our data versus the passive transfer of vaccine-derived sera. Importantly, as a result of the lack of available therapeutic options, and the previous successes with CP in the treatment of SARS-CoV and MERS-CoV, thousands of humans have been treated with CCP through compassionate use or clinical trial access channels where preliminary results suggest treatment is well tolerated (Joyner et al., 2020b). Meta-analysis of the available data suggest a clear benefit as measured by reduced hospitalization times and minimal risk where adverse events were classified as common to CP treatments in general rather than complications related specifically to CCP or ADE (Joyner et al., 2020a; Sun et al., 2020).

Ongoing clinical trials are working to refine the treatment approaches and our understanding of precisely what is needed to improve efficacy in terms of therapeutic dosing of CCP and issues related to safety (Joyner et al., 2020b). Several recent studies suggest that timing and dose may be critical to the success of treatment. For example, less therapeutic benefit was observed in patients that began treatment during advanced disease (Bradfute et al., 2020; Gharbharan et al., 2020). Further, a recent large scale clinical trial in India highlighted the importance of donor CCP potency where the median neutralizing antibody potency of donor CCP was 1:40 (interquartile range = 1:30-1:80) compared to our study wherein both treatment groups PRNT50 values were considerably higher (~1:128 low dose pool or 1:2048 high dose pool). The trial was terminated early as no clear therapeutic benefit was evident. (Agarwal et al., 2020).

Here we have provided detailed experimental evidence in a relevant animal model of COVID-19 which supports early administration of potent CCP. Given the exponential growth of a convalescent COVID-19 population, the availability of CCP as a treatment may serve as a therapeutic bridge until more potent and targeted immunotherapeutics such as monoclonal antibodies or other antiviral drugs become available.

## Acknowledgments

The authors would like to thank the UTMB Animal Resource Center for husbandry support of laboratory animals, Dr. Kevin Melody for assistance with animal studies, Dr. Liana Medina for assistance with processing BAL samples, and Kira Zapalac for assistance with clinical pathology and PCR assays. We also thank Drs. Luis Branco and Matt Boisen for generously providing the SARS-CoV2 anti-nucleoprotein and spike RBD ELISA assays as well as Dr. Thomas Ksiazek for the irradiated SARS-CoV2 whole cell lysate used in ELISA assays. The virus used in this publication was kindly provided by the European Virus Archive goes Global (EVAg) project that has received funding from the European Union’s Horizon 2020 research and innovation program under grant agreement No 653316.

## Author contributions

RWC and TWG conceived and designed the study. DJD, JBG, and TWG performed the SARS-CoV-2 challenge experiments. RWC, DJD, JBG, and TWG performed animal procedures and clinical observations. KNA and VB performed the clinical pathology assays. CW processed BAL samples. VB performed the SARS-CoV-2 infectivity assays. KNA performed the PCR. NSD performed the immunohistochemistry and *in situ* hybridization. CW performed ELISAs and multiplex assays. KAF performed the necropsies and analysis of the gross pathology, histopathology, immunohistochemistry and *in situ* hybridization. All authors analyzed the clinical pathology, virology, and immunology data. RWC, ANP, KAF, and TWG wrote the paper. All authors had access to all of the data and approved the final version of the manuscript.

## Data and materials availability

The data sets used and/or analyzed during the current study are available from the corresponding author on reasonable request.

## Materials and methods

### Virus

The virus (SARS-CoV-2/INMI1-Isolate/2020/Italy) was isolated on January 30, 2020 from the sputum of the first clinical case in Italy, a tourist visiting from the Hubei province of China that developed respiratory illness while traveling. The virus was initially passaged twice (P2) on Vero E6 cells; the supernatant and cell lysate were collected and clarified following a freeze/thaw cycle. This isolate is certified mycoplasma and Foot-and-Mouth Disease virus free. The complete sequence was submitted to GenBank (MT066156) and is available on the GISAID website (BetaCoV/Italy/INMI1-isl/2020: EPI_ISL_410545) upon registration. For *in vivo* challenge, the P2 virus was propagated on Vero E6 cells and the supernatant was collected and clarified by centrifugation making the virus used in this study a P3 stock.

### Animal challenge

Prior to enrollment in this study, AGMs (*Chlorocebus aethiops*; n=10; 6 males, 4 females; St Kitts origin, Worldwide Primates, Inc.) were tested for sero-reactivity to SARS-CoV-2. All animals were seronegative. Animals were anesthetized with ketamine and inoculated with a target dose of 5.0 x 10^5^ PFU of SARS-CoV-2 (SARS-CoV-2/INMI1-Isolate/2020/Italy) through combined intranasal (i.n.) and intratracheal (i.t.) inoculation, with the dose being divided evenly between both routes. All animals were longitudinally monitored for clinical signs of illness, respiration quality, and clinical pathology. All measurements requiring physical manipulation of the animals were performed under sedation by ketamine. Mucosal swabs were obtained using sterile swabs inserted into the mucosal cavity, gently rotated to maximize contact with the mucosal surface, and deposited into 2.0 mL screw-top tubes containing sterile MEM media supplemented to 10% with FBS.

### RNA isolation from SARS-CoV-2-infected AGMs

On specified procedure days (days 0, 2, 3, 4, 5), 100 μl of blood was added to 600 μl of AVL viral lysis buffer (Qiagen) for virus inactivation and RNA extraction. Following removal from the high containment laboratory, RNA was isolated from blood and swabs using the QIAamp viral RNA kit (Qiagen).

### Detection of SARS-CoV-2 load

RNA was isolated from blood and mucosal swabs and assessed using the CDC SARS-CoV-2 N2 assay primers/probe for reverse transcriptase quantitative PCR (RT-qPCR) [17]. SARS-CoV-2 RNA was detected using One-step probe RT-qPCR kits (Qiagen) run on the CFX96 detection system (Bio-Rad), with the following cycle conditions: 50°C for 10 minutes, 95°C for 10 seconds, and 45 cycles of 95°C for 10 seconds and 55°C for 30 seconds. Threshold cycle (*C*_*T*_) values representing SARS-CoV-2 genomes were analyzed with CFX Manager Software, and data are presented as GEq. To generate the GEq standard curve, RNA was extracted from supernatant derived from Vero E6 cells infected with SARS-CoV-2/INMI1-Isolate/2020/Italy was extracted and the number of genomes was calculated using Avogadro’s number and the molecular weight of the SARS-CoV-2 genome.

Infectious virus was quantitated by plaque assay on Vero E6 cells (ATCC CRL-1586) from all blood plasma and mucosal swabs, and bronchoalveolar lavage (BAL) samples. Briefly, increasing 10-fold dilutions of the samples were adsorbed to Vero E6 cell monolayers in duplicate wells (200μl). Cells were overlaid with EMEM medium plus 1.25% Avicel, incubated for 2 days, and plaques were counted after staining with 1% crystal violet in formalin. The limit of detection for this assay is 25 PFU/ml.

### Serum neutralization assay

Neutralization titers were calculated by determining the dilution of serum that reduced 50% of plaques (PRNT_50_). A standard 100 PFU amount of SARS-CoV-2 was incubated with two-fold serial dilutions of serum samples for one hour. The virus-serum mixture was then used to inoculate Vero E6 cells for 60 minutes. Cells were overlaid with EMEM medium plus 1.25% Avicel, incubated for 2 days, and plaques were counted after staining with 1% crystal violet in formalin.

### ELISA

SARS-CoV-2-specific IgG antibodies were measured in sera by ELISA at the indicated time points. Immunosorbent 96-well plates were coated overnight with each antigen. For total virus-specific IgG, plates were coated with a 1:1000 dilution of irradiated SARS-CoV-2 infected or normal Vero E6 lysate in PBS (pH 7.4) kindly provided by Dr. Thomas W. Ksiazek (UTMB). Nucleoprotein and Spike Receptor Binding Domain (RBD) ELISA kits were kindly provided by Zalgen Labs, LLC. Sera were initially diluted 1:100 and then two-fold through 1:51,200 in 4% BSA in 1× PBS or in Zalgen-provided reagents. After a one-hour incubation, plates were washed six times with wash buffer (1 x PBS with 0.2% Tween-20) and incubated for an hour with a 1:5000 dilution of horseradish peroxidase (HRP)-conjugated anti-primate IgG antibody (Fitzgerald Industries International; Cat: 43R-IG020HRP). RT SigmaFast O-phenylenediamine (OPD) substrate (P9187, Sigma) was added to the wells after six additional washes to develop the colorimetric reaction. The reaction was stopped with 3M sulfuric acid 5-10 minutes after OPD addition and absorbance values were measured at a wavelength of 492nm on a spectrophotometer (Biotek Cytation5 system). Absorbance values were normalized by subtracting uncoated wells from antigen-coated wells at the corresponding serum dilution. End-point titers were defined as the reciprocal of the last adjusted serum dilution with a value ≥ 0.20.

### Histopathology and immunohistochemistry

Necropsy was performed on all subjects euthanized at 5 dpi. Tissue samples of all major organs were collected for histopathologic and immunohistochemical (IHC) examination and were immersion-fixed in 10% neutral buffered formalin for > 7 days. Specimens were processed and embedded in paraffin and sectioned at 5 μm thickness. For IHC, specific anti-SARS immunoreactivity was detected using an anti-SARS nucleocapsid protein rabbit primary antibody at a 1:800 dilution for 60 minutes (Novusbio). The tissue sections were processed for IHC using the ThermoFisher Scientific Lab Vision Autostainer 360 (ThermoFisher Scientific). Secondary antibody used was biotinylated goat anti-rabbit IgG (Vector Laboratories) at 1:200 for 30 minutes followed by Vector Streptavidin Alkaline Phosphatase at a dilution of 1:200 for 20 min (Vector Laboratories). Slides were developed with Bio-Red (Biopath) for 7 minutes and counterstained with hematoxylin for one minute. For IHC, specific anti-fibrin was detected using an anti-fibrin monoclonal mouse primary antibody at a 1:3200 dilution for 60 minutes (Sekisui Diagnostics). The tissue sections were processed for IHC using the ThermoFisher Scientific Lab Vision Autostainer 360 (ThermoFisher Scientific). Secondary antibody used was biotinylated goat anti-mouse IgG (Vector Laboratories) at 1:200 for 30 minutes followed by Vector Streptavidin Alkaline Phosphatase at a dilution of 1:200 for 20 min (Vector Laboratories). Slides were developed with Bio-Red (Biopath Laboratories) for 7 minutes and counterstained with hematoxylin for one minute. Tissues were stained following package instructions for collagen with the Trichrome One-Step Blue & Red Stain Kit (American MasterTech Scientific Laboratory Supplies).

### Bead-based cytokine immunoassays

Concentrations of immune mediators were determined by flow cytometry using LegendPlex multiplex technology (BioLegend). Serum levels of cytokines/chemokines were quantified using Nonhuman Primate Inflammation 13-plex (1:4 dilution) kit. Samples were processed in duplicate following the kit instructions and recommendations. Following bead staining and washing, 4000 bead events were collected on a FACS Canto II cytometer (BD Biosciences) using BD FACS Diva software. The raw .fcs files were analyzed with BioLegend’s cloud-based LEGENDplex™ Data Analysis Software.

### Statistical Analysis

The data was analyzed and graphed in GraphPad Prism 8.0. Lesion severity scores and tissue PCR and plaque assay titers were analyzed using either a one-way ANOVA or a two-way ANOVA supported by Tukey’s multiple comparisons test. ELISAs and LegendPlex assays were also analyzed using a two-way ANOVA and Tukey’s multiple comparisons test. No data points were excluded for our analyses. Significance cut-off values are outlined per data in their respective results sections only where significant differences in groups compared existed.

---

**Supplemental Figure S1:**
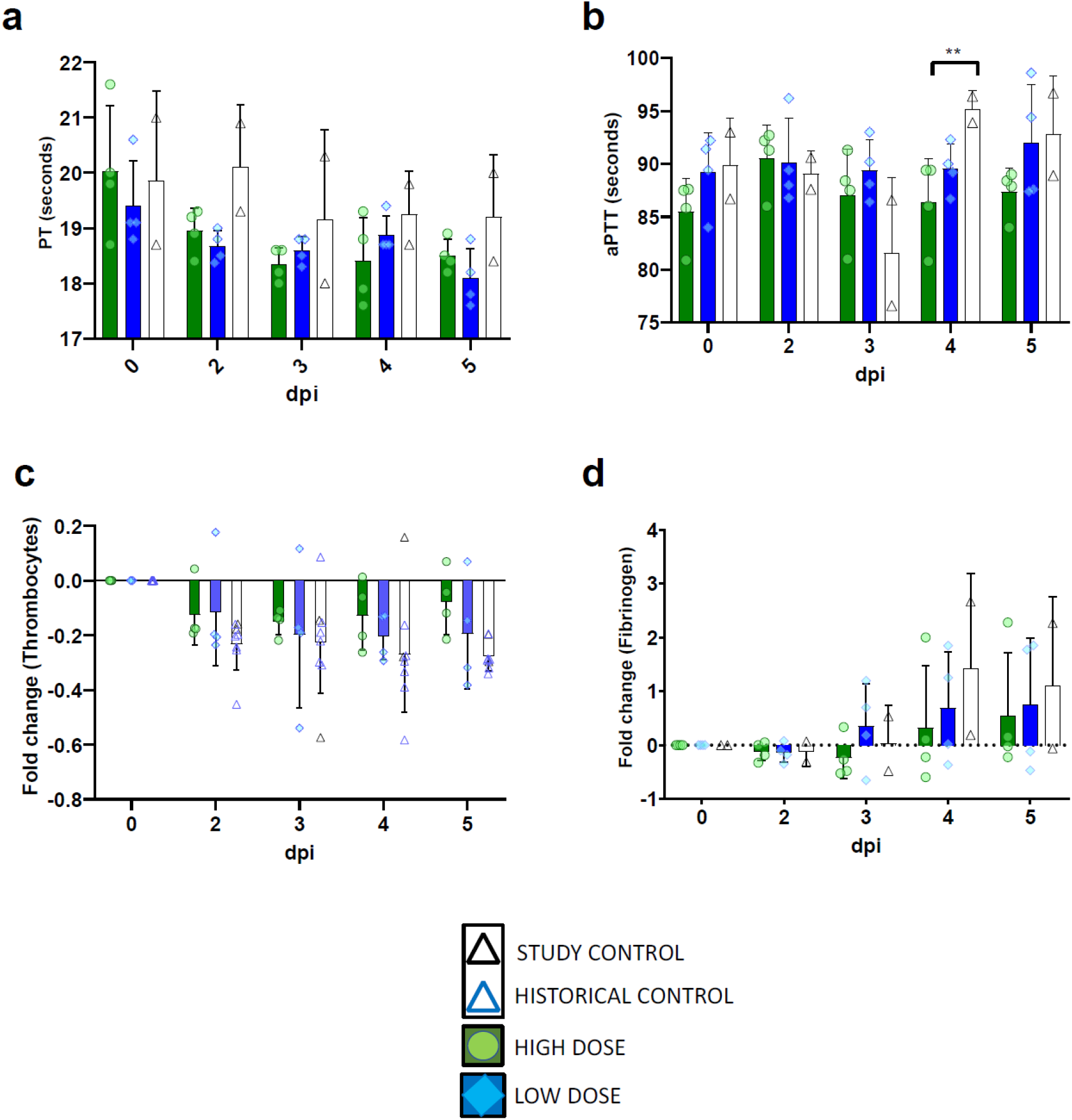
Coagulation profiles of AGMs following challenge with SARS-CoV-2 and treatment with convalescent sera. Prothrombin time (PT) **(a)**, activated partial thromboplastin time (aPTT) **(b)**, and fibrinogen levels (shown as fold change from individual subject baseline values) (b), were measured using a Vetscan VSPro coagulation analyzer. Note that historical control animals were not included for analysis in panels **(a-d)** due to a lack of available data for these parameters. Bars indicate the mean for each cohort at the indicated timepoint, values for individual animals within each cohort are shown as color-coded symbols. Error bars indicate the upper bound SD. Statistical significance was assessed by two-way ANOVA with the Geisser-Greenhouse correction without the assumption of sphericity, followed by Tukey’s post-hoc test for multiple comparisons to assess differences between cohorts at matched time points. Asterisk indicates significance (0.05 ≤ p ≤ 0.01). (**d**) Fold change from individual subject baselines (0 dpi) in absolute counts of thrombocytes were calculated for each subject at the indicated timepoints **(A)**. Historical control animals from a previous study (Woolsey et al., 2020) utilizing the homologous virus were included for statistical purposes.

## References

Agarwal, A., Mukherjee, A., Kumar, G., Chatterjee, P., Bhatnagar, T., Malhotra, P., Latha, B., Bundas, S., Kumar, V., Dosi, R., et al. (2020). Convalescent plasma in the management of moderate COVID-19 in India: An open-label parallel-arm phase II multicentre randomized controlled trial (PLACID Trial). medRxiv, 2020.2009.2003.20187252.

Arabi, Y., Balkhy, H., Hajeer, A.H., Bouchama, A., Hayden, F.G., Al-Omari, A., Al-Hameed, F.M., Taha, Y., Shindo, N., Whitehead, J., et al. (2015). Feasibility, safety, clinical, and laboratory effects of convalescent plasma therapy for patients with Middle East respiratory syndrome coronavirus infection: a study protocol. SpringerPlus. 4, 709.

Beigel, J.H., Tomashek, K.M., Dodd, L.E., Mehta, A.K., Zingman, B.S., Kalil, A.C., Hohmann, E., Chu, H.Y., Luetkemeyer, A., Kline, S., et al. (2020). Remdesivir for the Treatment of Covid-19 — Preliminary Report. New England Journal of Medicine.

Bradfute, S.B., Hurwitz, I., Yingling, A.V., Ye, C., Cheng, Q., Noonan, T.P., Raval, J.S., Sosa, N.R., Mertz, G.J., Perkins, D.J., et al. (2020). Severe Acute Respiratory Syndrome Coronavirus 2 Neutralizing Antibody Titers in Convalescent Plasma and Recipients in New Mexico: An Open Treatment Study in Patients With Coronavirus Disease 2019. The Journal of infectious diseases.

Casadevall, A., Dadachova, E., and Pirofski, L.A. (2004). Passive antibody therapy for infectious diseases. Nature reviews Microbiology. 2, 695–703.

CCPP19 (2020). National COVID-19 Convalescent Plasma Project-Component 3: Clinical Trials.

Chan, J.F.-W., Zhang, A.J., Yuan, S., Poon, V.K.-M., Chan, C.C.-S., Lee, A.C.-Y., Chan, W.-M., Fan, Z., Tsoi, H.-W., Wen, L., et al. (2020). Simulation of the clinical and pathological manifestations of Coronavirus Disease 2019 (COVID-19) in golden Syrian hamster model: implications for disease pathogenesis and transmissibility. Clinical Infectious Diseases.

Corti, D., Passini, N., Lanzavecchia, A., and Zambon, M. (2016). Rapid generation of a human monoclonal antibody to combat Middle East respiratory syndrome. Journal of Infection and Public Health. 9, 231–235.

Cross, R.W., Agans, K.N., Prasad, A.N., Borisevich, V., Woolsey, C., Deer, D.J., Dobias, N.S., Geisbert, J.B., Fenton, K.A., and Geisbert, T.W. (2020). Intranasal exposure of African green monkeys to SARS-CoV-2 results in acute phase pneumonia with shedding and lung injury still present in the early convalescence phase. Virol J. 17, 125–125.

Gharbharan, A., Jordans, C.C.E., GeurtsvanKessel, C., den Hollander, J.G., Karim, F., Mollema, F.P.N., Stalenhoef, J.E., Dofferhoff, A., Ludwig, I., Koster, A., et al. (2020). Convalescent Plasma for COVID-19. A randomized clinical trial. medRxiv. 2020.2007.2001.20139857.

Hartman, A.L., Nambulli, S., McMillen, C.M., White, A.G., Tilston-Lunel, N.L., Albe, J.R., Cottle, E., Dunn, M.D., Frye, L.J., Gilliland, T.H., et al. (2020). SARS-CoV-2 infection of African green monkeys results in mild respiratory disease discernible by PET/CT imaging and shedding of infectious virus from both respiratory and gastrointestinal tracts. PLOS Pathogens. 16, e1008903.

Imai, M., Iwatsuki-Horimoto, K., Hatta, M., Loeber, S., Halfmann, P.J., Nakajima, N., Watanabe, T., Ujie, M., Takahashi, K., Ito, M., et al. (2020). Syrian hamsters as a small animal model for SARS-CoV-2 infection and countermeasure development. Proceedings of the National Academy of Sciences. 117, 16587.

Jackson, L.A., Anderson, E.J., Rouphael, N.G., Roberts, P.C., Makhene, M., Coler, R.N., McCullough, M.P., Chappell, J.D., Denison, M.R., Stevens, L.J., et al. (2020). An mRNA Vaccine against SARS-CoV-2 — Preliminary Report. New England Journal of Medicine.

Johnston, S.C., Jay, A., Raymond, J.L., Rossi, F., Zeng, X., Scruggs, J., Dyer, D., Frick, O., Moore, J., Berrier, K., et al. (2020). Development of a Coronavirus Disease 2019 Nonhuman Primate Model Using Airborne Exposure. bioRxiv. 2020.2006.2026.174128.

Joyner, M.J., Klassen, S.A., Senefeld, J., Johnson, P.W., Carter, R.E., Wiggins, C.C., Shoham, S., Grossman, B.J., Henderson, J.P., Musser, J.M., et al. (2020a). Evidence favouring the efficacy of convalescent plasma for COVID-19 therapy. medRxiv. 2020.2007.2029.20162917.

Joyner, M.J., Wright, R.S., Fairweather, D., Senefeld, J.W., Bruno, K.A., Klassen, S.A., Carter, R.E., Klompas, A.M., Wiggins, C.C., Shepherd, J.R.A., et al. (2020b). Early safety indicators of COVID-19 convalescent plasma in 5000 patients. The Journal of Clinical Investigation 130, 4791–4797.

Laing, A.G., Lorenc, A., del Molino del Barrio, I., Das, A., Fish, M., Monin, L., Muñoz-Ruiz, M., McKenzie, D.R., Hayday, T.S., Francos-Quijorna, I., et al. (2020). A dynamic COVID-19 immune signature includes associations with poor prognosis. Nature Medicine.

Liu, L., Wei, Q., Lin, Q., Fang, J., Wang, H., Kwok, H., Tang, H., Nishiura, K., Peng, J., Tan, Z., et al. (2019). Anti-spike IgG causes severe acute lung injury by skewing macrophage responses during acute SARS-CoV infection. JCI Insight. 4.

Lu, S., Zhao, Y., Yu, W., Yang, Y., Gao, J., Wang, J., Kuang, D., Yang, M., Yang, J., Ma, C., et al. (2020). Comparison of nonhuman primates identified the suitable model for COVID-19. Signal Transduction and Targeted Therapy 5, 157.

Mair-Jenkins, J., Saavedra-Campos, M., Baillie, J.K., Cleary, P., Khaw, F.M., Lim, W.S., Makki, S., Rooney, K.D., Nguyen-Van-Tam, J.S., and Beck, C.R. (2015). The effectiveness of convalescent plasma and hyperimmune immunoglobulin for the treatment of severe acute respiratory infections of viral etiology: a systematic review and exploratory meta-analysis. The Journal of infectious diseases. 211, 80–90.

Malani, A.N., Sherbeck, J.P., and Malani, P.N. (2020). Convalescent Plasma and COVID-19. JAMA 324, 524–524.

Rockx, B., Kuiken, T., Herfst, S., Bestebroer, T., Lamers, M.M., Oude Munnink, B.B., De Meulder, D., van Amerongen, G., van de Brand, J., Okba, N.M.A., et al. (2020). Comparative pathogenesis of COVID-19, MERS, and SARS in a nonhuman primate model. Science. 368, 1012.

Sariol, A., and Perlman, S. (2020). Lessons for COVID-19 Immunity from Other Coronavirus Infections. Immunity. 53, 248–263.

Shen, C., Wang, Z., Zhao, F., Yang, Y., Li, J., Yuan, J., Wang, F., Li, D., Yang, M., Xing, L., et al. (2020). Treatment of 5 Critically Ill Patients With COVID-19 With Convalescent Plasma. JAMA 323, 1582–1589.

Singh, D.K., Ganatra, S.R., Singh, B., Cole, J., Alfson, K.J., Clemmons, E., Gazi, M., Gonzalez, O., Escobedo, R., Lee, T.-H., et al. (2020). SARS-CoV-2 infection leads to acute infection with dynamic cellular and inflammatory flux in the lung that varies across nonhuman primate species. bioRxiv 2020.2006.2005.136481.

Speranza, E., Williamson, B.N., Feldmann, F., Sturdevant, G.L., Pérez-Pérez, L., Mead-White, K., Smith, B.J., Lovaglio, J., Martens, C., Munster, V.J., et al. (2020). SARS-CoV-2 infection dynamics in lungs of African green monkeys. bioRxiv 2020.2008.2020.258087.

Sun, M., Xu, Y., He, H., Zhang, L., Wang, X., Qiu, Q., Sun, C., Guo, Y., Qiu, S., and Ma, K. (2020). A potentially effective treatment for COVID-19: A systematic review and meta-analysis of convalescent plasma therapy in treating severe infectious disease. Int J Infect Dis. 98, 334–346.

US-FDA (2020). FDA Issues Emergency Use Authorization for Convalescent Plasma as Potential Promising COVID–19 Treatment, Another Achievement in Administration’s Fight Against Pandemic.

van Doremalen, N., Falzarano, D., Ying, T., de Wit, E., Bushmaker, T., Feldmann, F., Okumura, A., Wang, Y., Scott, D.P., Hanley, P.W., et al. (2017). Efficacy of antibody-based therapies against Middle East respiratory syndrome coronavirus (MERS-CoV) in common marmosets. Antiviral research 143, 30–37.

Woolsey, C., Borisevich, V., Prasad, A.N., Agans, K.N., Deer, D.J., Dobias, N.S., Heymann, J.C., Foster, S.L., Levine, C.B., Medina, L., et al. (2020). Establishment of an African green monkey model for COVID-19. bioRxiv : the preprint server for biology. 2020.2005.2017.100289.

World.Health.Organisation (2020). Coronavirus Disease 2019.

Yeh, K.-M., Chiueh, T.-S., Siu, L.K., Lin, J.-C., Chan, P.K.S., Peng, M.-Y., Wan, H.-L., Chen, J.-H., Hu, B.-S., Perng, C.-L., et al. (2005). Experience of using convalescent plasma for severe acute respiratory syndrome among healthcare workers in a Taiwan hospital. Journal of Antimicrobial Chemotherapy. 56, 919–922.

Zhu, F.-C., Guan, X.-H., Li, Y.-H., Huang, J.-Y., Jiang, T., Hou, L.-H., Li, J.-X., Yang, B.-F., Wang, L., Wang, W.-J., et al. (2020). Immunogenicity and safety of a recombinant adenovirus type-5-vectored COVID-19 vaccine in healthy adults aged 18 years or older: a randomised, double-blind, placebo-controlled, phase 2 trial. The Lancet. 396, 479–488.

